# DPPA2 and DPPA4 are necessary to establish a totipotent state in mouse embryonic stem cells

**DOI:** 10.1101/447755

**Authors:** Alberto De Iaco, Alexandre Coudray, Julien Duc, Didier Trono

## Abstract

After fertilization of the transcriptionally silent oocyte, expression from both parental chromosomes is launched through so-called zygotic genome activation (ZGA), occurring in the mouse at the 2-cell stage. Amongst the first elements to be transcribed are the *Dux* gene, the product of which secondarily induces a wide array of ZGA genes, and a subset of evolutionary recent LINE-1 retrotransposons, which regulate chromatin accessibility in the early embryo. The maternally-inherited factors that activate *Dux* and LINE-1 transcription have so far remained unknown. Here we identify the paralog proteins DPPA2 and DPPA4 as responsible for this process.

## Introduction

Mammalian embryonic development begins with the fertilization of a transcriptionally silent oocyte by the spermatozoa, resulting in the formation of a totipotent zygote. Transcription from the two parental genomes ensues through a phenomenon known as zygotic or embryonic genome activation (ZGA or EGA). Murine ZGA occurs at the 2-cell stage, and members of the DUX family of transcription factors were recently identified as important inducers of early embryonic genes in mouse, human and likely all placental mammals (De Iaco et al. 2017; Hendrickson et al. 2017; Whiddon et al. 2017). However, murine and human DUX are expressed only after fertilization of the oocytes. Moreover, a subset of LINE-1 retrotransposons is transiently activated during this period, a DUX-independent phenomenon important for regulating chromatin accessibility in the murine pre-implantation embryo (Jachowicz et al. 2017). Maternally inherited factors are thus likely responsible for inducing the transcription of both *Dux* and LINE-1 in the nascent embryo. *Dppa2* (developmental pluripotency-associated 2) and *Dppa4* (developmental pluripotency-associated 4) are paralog genes conserved in mammals, and found as unique *Dppa2* orthologs in amphibians, reptiles and marsupials (Siegel et al. 2009). All *Dppa2* and *Dppa4* ortholog genes have a conserved SAP (SAF-A/B, Acinus and PIAS) motif, important for DNA-binding, and a C-terminal domain of unknown function (Aravind and Koonin 2000). These genes were first described for their expression profile restricted to pluripotent cells and the germline (Bortvin et al. 2003; Maldonado-Saldivia et al. 2007). It was subsequently demonstrated that DPPA2 and DPPA4 activate transcription of genes important for germ cell development in mouse embryonic stem cells (mESCs), and are both essential for murine embryogenesis (Madan et al. 2009; Nakamura et al. 2011). Embryonic development is impaired when DPPA4 is depleted from murine oocytes, suggesting that the maternally inherited fraction of the protein plays an important role in the early pre-implantation period (Madan et al. 2009). How DPPA2 and DPPA4 activate transcription of target genes is unclear, but both proteins associate with transcriptionally active chromatin in mESCs (Masaki et al. 2007; Engelen et al. 2015), and promoters of DPPA2-stimulated genes acquire repressive epigenetic marks upon loss of this factor in mESCs (Nakamura et al. 2011). This suggests that DPPA2 and DPPA4 act as epigenetic modifiers. In this study, we demonstrate that DPPA2 and DPPA4 act as inducers of *Dux* and LINE1 transcription, thus promoting the establishment of a totipotent state.

## Results

### DPPA2 and DPPA4 are expressed prior to ZGA

The downregulation of genes expressed in murine 2C embryos when *Dppa4* is deleted in mESCs (Madan et al. 2009) suggested that its product might regulate ZGA. We thus examined the expression of this gene and its paralog *Dppa2* in pre-implantation embryos. We found *Dppa2* transcript levels to be high from oocytes to blastocysts, and their *Dppa4* counterparts to raise only at ZGA (Figure 1A, Supplementary Figure 1A) (Kobayashi et al. 2012; Deng et al. 2014). Therefore, both DPPA2 and DPPA4 are present when *Dux*, a strong inducer of ZGA, starts being transcribed.

*Dppa4* encodes for two protein isoforms, one full-length and one lacking the SAP domain (DPPA4ΔSAP) (Figure 1B, Supplementary Figure 1B) (Madan et al. 2009). The SAP domain promotes nuclear localization, suggesting that the truncated DPPA4 variant is less prone to associate with chromatin (Madan et al. 2009). Interestingly, we detected only low levels of full-length *Dppa4* transcripts in zygotes and 2C embryos, likely the remnants of maternal transcripts, while their truncated counterparts were highly expressed in middle and late 2C, that is, upon ZGA (Figure 1C). This indicates that maternally-inherited and ZGA-produced DPPA4 isoforms are different, and may have distinct functions. In mESCs, we detected only full-length *Dppa4* transcripts (Supplementary figure 1C), and their levels were comparable in *wild-type* (WT) and *Dux* KO cells, indicating that *Dppa4* expression is not regulated by DUX (Supplementary Figure 1D).

**Figure 1.**
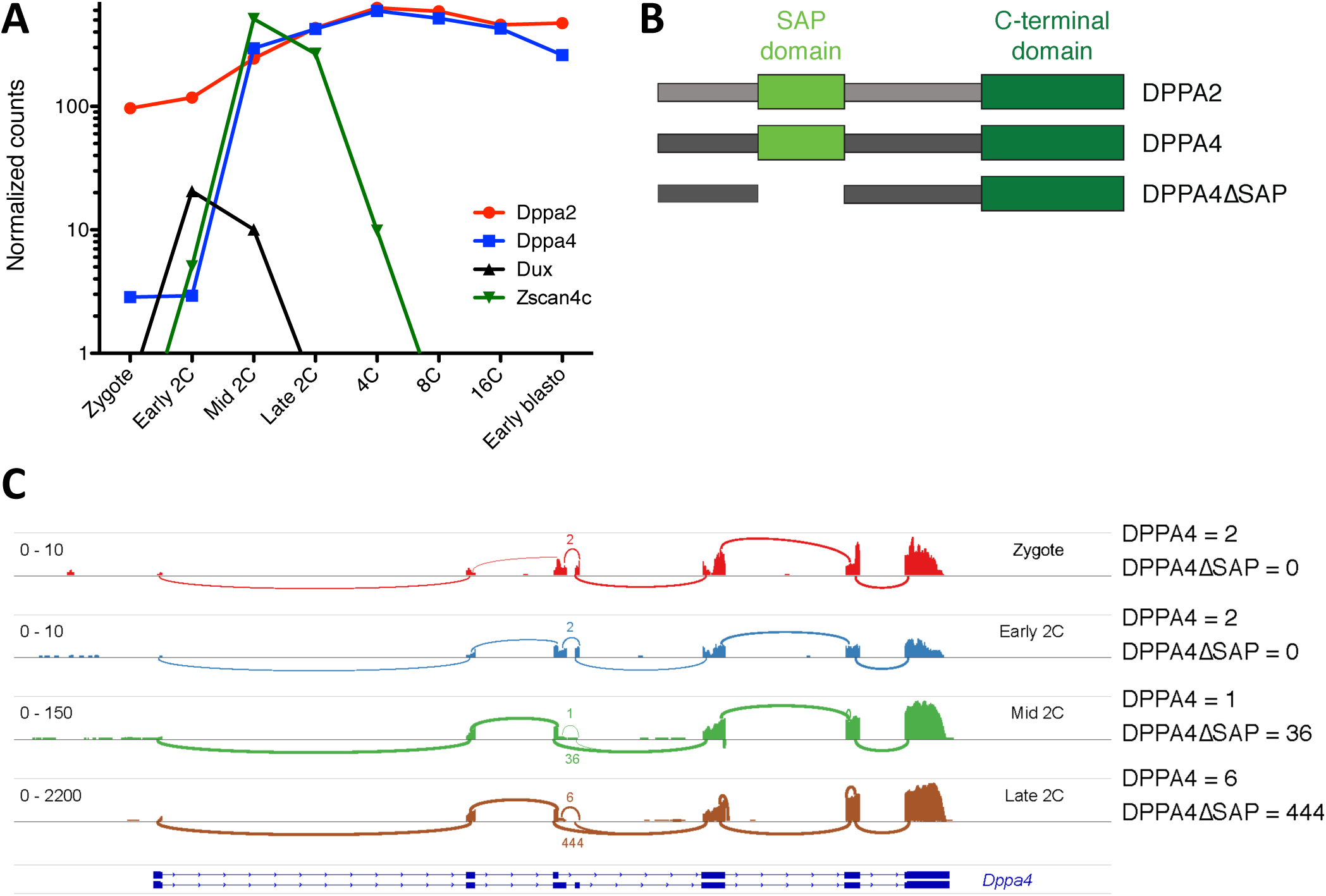
DPPA2 and DPPA4 are expressed throughout pre-implantation embryo development. (**A**) Comparative expression during murine pre-implantation embryo development of *Dppa2* (red), *Dppa4* (blue), *Dux* (black), and *Zscan4c* (ZGA marker, green). Each dot represents the average value of single-cell RNA-seq from (Deng et al. 2014) (**B**) Schematic representation of DPPA2, DPPA4 full-length and DPPA4 truncated proteins. The conserved domains are shown: SAP (light green), C-terminal (dark green). (**C**) Sashimi plot representing coverage on the *Dppa4* gene of an RNA sequencing analysis of murine zygotes, early 2C, mid 2C and late 2C (A). Arcs display the number of reads split across junctions (splicing events). Numbers represent the number of splicing events detected at a junction.

### DPPA2 and DPPA4 nduce expression of DUX and other ZGA genes in 2C-like mESCs

A subpopulation of mESCs known as 2C-like cells cycles through a totipotent-like state where *Dux* and its ZGA-specific target genes are transiently expressed (Macfarlan et al. 2012; De Iaco et al. 2017). By analyzing a publicly available RNA sequencing (RNA-seq) dataset from mESCs sorted for expression of a 2C-specific reporter system (MERVL-GFP, *Zscan4*-Tomato) (Eckersley-Maslin et al. 2016), we found higher level of *Dppa2* but not *Dppa4* transcripts in this subpopulation (Figure 2A, Supplementary Figure 2A). To ask whether DPPA2 and DPPA4 are necessary for transition through this 2C-like state, we transduced mESCs carrying the MERVL-GFP reporter, the expression of which we previously demonstrated to be DUX-dependent (De Iaco et al. 2017), with lentiviral vectors expressing shRNAs directed against *Dppa2* or *Dppa4* (Supplementary Figure 2BC). We found that expression of *Dux*, its downstream target *Zscan4*, and the MERVL-GFP reporter was lost upon DPPA2 or DPPA4 depletion. To confirm these results, we deleted *Dppa2* or *Dppa4* from mESCs by CRISPR/Cas9-mediated genome editing (Figure 2B). Transcriptome analyses revealed a complete loss of transcripts from *Dux* and most of its previously defined downstream target genes (Figure 2CD, Supplementary Figure 2D). Expression of some of these genes (*Zscan4* and *Tdpoz4*) was rescued when *Dux* was ectopically expressed in *Dppa2* or *Dppa4* KO mESCs (Figure 2E), suggesting that DPPA2 and DPPA4 act upstream of DUX in the establishment of a 2C-like state in mESCs. We also identified another subset of 1155 genes downregulated in both *Dppa2* and *Dppa4* KO mESCs that were not controlled by DUX, including *Mael*, *Tdrd1* and *Prex2* (Figure 2F, Supplementary Figure 2EF). These genes were more expressed in 2C-like cells than in the rest of mESCs population (Figure 2G). Nevertheless, only 1/10 of these genes were transcribed specifically at ZGA (Supplementary Figure 2G), pointing to differences in the transcriptomes of 2C embryos and 2C-like ES cells (Madan et al. 2009; Nakamura et al. 2011). We overexpressed hemagglutinin (HA)-tagged forms of DPPA2 or DPPA4 in the corresponding KO mESCs (Figure 3A). Expression of *Mael*, *Tdrd1* and *Prex2* was restored, but not that of *Dux* and downstream targets such as *Zscan4* or *Tdpoz4*, perhaps because levels of complementation achieved for either DPPA2 or DPPA4 proteins were below those detected for their endogenous counterparts in control cells (Supplementary Figure 3A).

**Figure 2.**
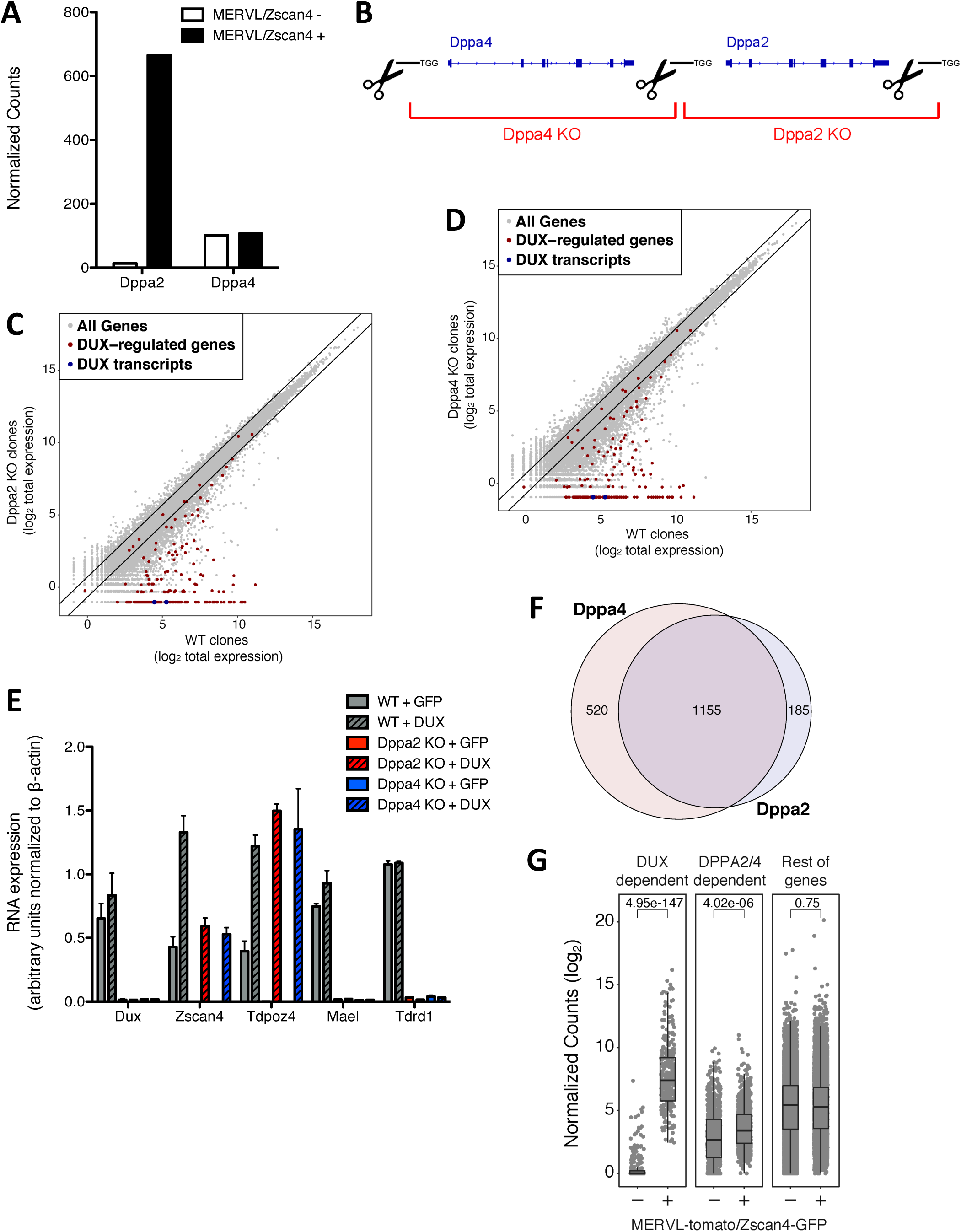
DPPA2 and DPPA4 regulate expression of *Dux* in mESCs. (**A**) Average expression of *Dppa2* and *Dppa4* in a single-cell RNA-seq analysis of mESCs sorted for expression of both Tomato and GFP reporters driven by MERVL and *Zscan4* promoters, respectively, and the double-negative population. (**B**) Schematic representation of the CRISPR/Cas9 approach to deplete DPPA2 or DPPA4 from mESCs. Black lines represent the sgRNAs used. RNA-seq analysis of WT and *Dppa2* (**C**) or Dppa4 (**D**) KO mESC clones. The dot plot displays the average gene expression of three independent clones from each cell type. (**E**) Comparative expression by qPCR of *Dux*, two downstream targets of DUX (*Zscan4* and *Tdpoz4*), and two DUX-independent targets of DPPA2 and DPPA4 (*Mael* and *Tdrd1*) in mESCs depleted of endogenous DPPA2 and DPPA4 and overexpressing ectopically DUX or GFP as a control. Expression was normalized to *Actb*. n = 3 (**F**) Venn diagram representing the overlap of genes dowregulated in *Dppa2* and *Dppa4* KO compared to WT cells. (**G**) Comparative expression of DUX-dependent, DPPA2 and DPPA4-dependent, and the rest of the genes in mESCs sorted for expression of both Tomato and GFP reporters driven by MERVL and *Zscan4* promoters, respectively, and the double-negative population (unpaired t-test).

### DPPA2 and DPPA4 bind the promoters of *Dux* and other ZGA genes

In spite of this only partial complementation, chromatin immunoprecipitation studies revealed that both DPPA2 and DPPA4 associated with the 5’ region of genes downregulated in their absence including *Dux*, and that these factors were not enriched at DUX-recruiting promoters (Figure 3BC, Supplementary Figure 3B). This strongly suggests that DPPA2 and DPPA4 are direct activators of *Dux*, and secondarily induce genes such as *Zscan4* and other DUX-driven targets. Interestingly, DPPA4 lacking the SAP domain was still able to bind the 5’ end of *Dux*, albeit less strongly than the full-length protein, and also associated with a sizeable fraction of DPPA4-controlled genes, the expression of which it could partly rescue by complementation of *Dppa4* KO mESCs (Supplementary Figure 3C). Therefore, the SAP domain is absolutely essential neither for the genomic recruitment nor for the transactivating activity of DPPA4. Finally, DPPA2 and DPPA4 were less enriched at the promoter of *Dux*, *Mael* and *Tdrd1* in mESCs deleted for the reciprocal paralog, compared with control cells (Supplementary Figure 3D). It indicates that DPPA2 and DPPA4 are best recruited jointly at target genes, consistent with the recent demonstration of their heterodimerization potential (Hernandez et al. 2018). We attempted to delineate more precisely the sequence motifs recognized by DPPA2 and DPPA4, and could only determine that the two proteins favor GC-rich regions such as found at the 5’ end of *Dux* or CpG islands (Figure 3D, Supplementary Figure 3EF).

**Figure 3.**
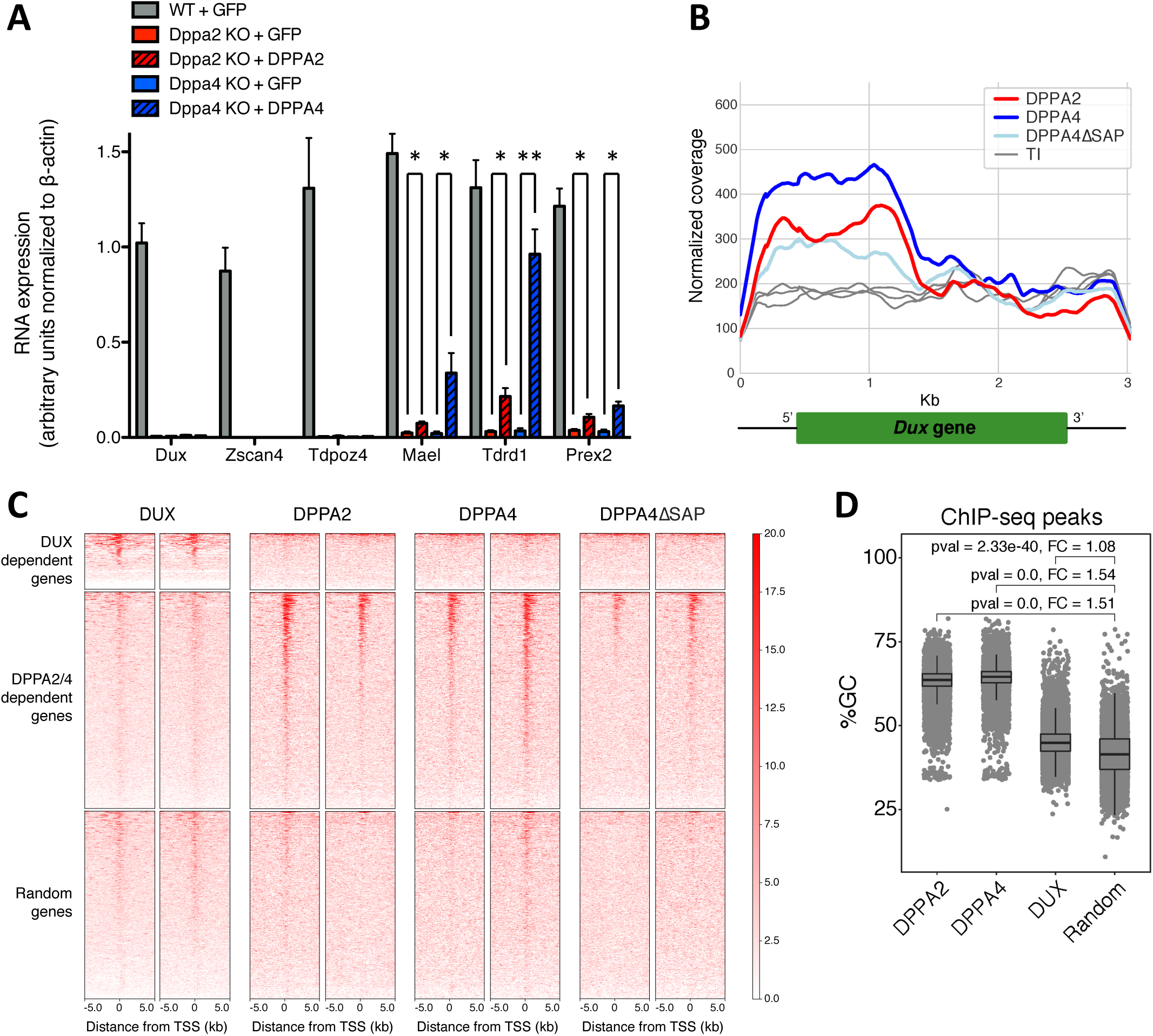
DPPA2 and DPPA4 associate with the promoter of their target genes in mESCs. (**A**) Comparative expression by qPCR of *Dux*, two downstream targets of DUX (*Zscan4* and *Tdpoz4*), and three targets of DPPA2 and DPPA4 independent of DUX (*Mael, Tdrd1, Prex2*) in mESCs depleted of endogenous DPPA2 and DPPA4 and overexpressing ectopically DPPA2, DPPA4 or GFP as a control (* p < 0.05, ** p < 0.01, unpaired t-test, n = 3). (**B**) Average coverage normalized for sequencing depth of the ChIP-seq signal of DPPA2, DPPA4 or DPPA4 truncated of the SAP domain overexpressed in mESCs in a window of 500bp around the *Dux* gene (n=2). Total input (TI) is shown in gray. Peaks over the *Dux* gene were called in DPPA2, DPPA4 and DPPA4ΔSAP ChIP-seq. (**C**) Heatmap showing the distribution of DUX, DPPA2, DPPA4 and DPPA4ΔSAP coverage in a ± 5kb window around the TSS of DUX-dependent, DPPA2 and DPPA4-dependent and random genes in mESCs. (**D**) Boxplot/Jitterplot representing the percentage of GC nucleotide contents in the peaks from DPPA2, DPPA4 and DUX ChIP, and in a random shuffle of peaks.

### DPPA2 and DPPA4 regulate the expression of young LINE-1s in mESCs

Upon comparing the transposcriptomes (sum of transposable elements-derived transcripts) of WT, *Dppa2-* and *Dppa4*-deleted mESCs, we found that levels of MERVL-int, L1Md_T, and L1Md_A RNAs dramatically dropped in the KO cells (Figure 4AB, Supplementary Figure 4A). We further determined that, similar to MERVL, L1Md_T and L1Md_A were more highly expressed in the 2C-like subpopulation of mESCs (Figure 4C).

**Figure 4.**
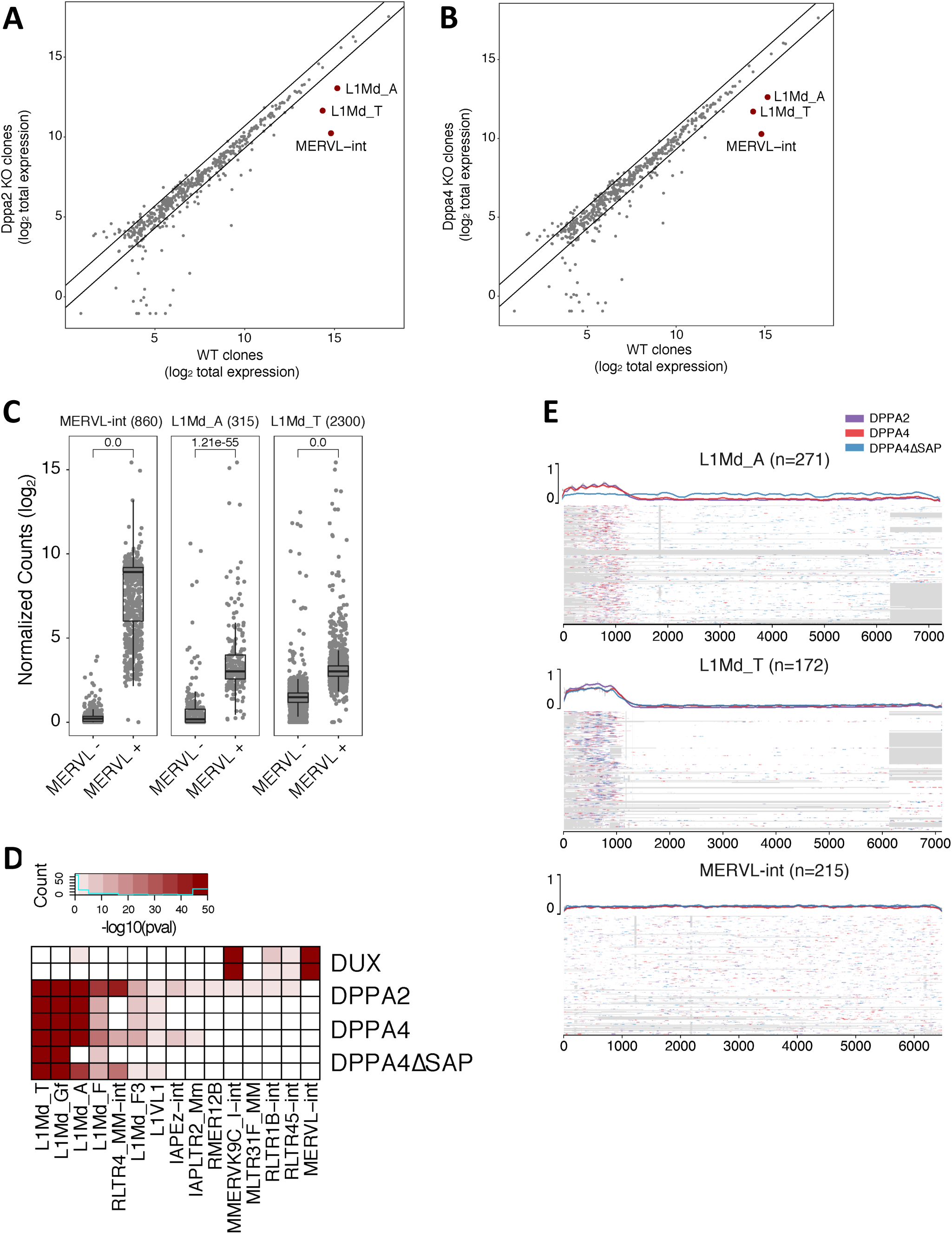
DPPA2 and DPPA4 directly regulate expression of young LINE-1 families in mESCs. Total expression of transposable element subfamilies in (**A**) *Dppa2* KO or (**B**) *Dppa4* KO compared to WT mESC clones. (**C**) RNA-seq analysis of MERVL-int, L1Md_A and L1Md_T in mESCs sorted for expression of both Tomato and GFP reporters driven by MERVL and *Zscan4* promoters, respectively, and the double-negative population. (**D**) Heatmap displaying the enrichment of DUX, DPPA2, DPPA4 and DPPA4ΔSAP binding at different transposable element families. Every subfamily enriched (pval < 0.05) in at least one DPPA2, DPPA4 or DPPA4ΔSAP replicate are shown. (**E**) MSA plot of DPPA2, DPPA4 and DPPA4ΔSAP ChIP-seq in mESCs showing enrichment over L1Md_A, L1Md_T and MERVL-int sequences. Gaps are in grey and sequences in white with overlap of color-coded ChIP-seq signals. Upper plot shows average coverage of the signal over the aligned TEs.

MERVL is an endogenous retrovirus previously identified as a direct target of DUX, but L1Md_T, and L1Md_A are two evolutionarily recent subfamilies of murine LINE-1 elements, the expression of which is DUX-independent (Sookdeo et al. 2013; Whiddon et al. 2017). Accordingly, in *Dux* KO mESCs, expression of MERVL was completely lost but that of L1Md_T and L1Md_A only slightly reduced (Supplementary Figure 4B). However, L1Md_T and L1Md_A transcript levels dropped when DPPA2 and DPPA4 were depleted from *Dux* KO mESCs by RNA interference (Supplementary Figure 4C). Furthermore, chromatin immunoprecipitation analyses found HA-tagged forms of DPPA2 and DPPA4 enriched at the 5’-end of L1Md_T and L1Md_A, but not MERVL-int (Figure 4DE). Interestingly, the 5’ untranslated region of LINE-1 responsible for recruiting DPPA2 and DPPA4 has a high GC content (Dai et al. 2017). Of note, deleting the SAP domain of DPPA4 reduced its recruitment at L1Md_A but not L1Md_T integrants (Figure 4DE).

## Discussion

This work identifies DPPA2 and DPPA4 as activators of *Dux* and LINE-1 retrotransposons induced during murine ZGA and in 2C-like mESCs. The presence of DPPA2 and DPPA4 at fertilization and the partial requirement of maternal DPPA4 further suggest that the two proteins promote the establishment of the totipotent early embryo (Madan et al. 2009). We found high levels of *Dppa2* throughout pre-implantation development, while two different isoforms of *Dppa4* were detected: low levels of the full-length form in oocytes and zygotes, and a form lacking the SAP domain expressed from ZGA to blastocyst. While the SAP domain of DPPA4 was found to foster its nuclear localization (Madan et al. 2009), we observed that DPPA4ΔSAP was still able to bind the promoters and activate expression of some of the gene and transposon targets of the full-length protein, albeit with reduced efficiency. Still, presence of distinct DPPA4 isoforms in the zygote and after cleavage suggests that they fulfill different functions in the totipotent cell and at later stages of pre-implantation development. In *Xenopus*, gastrulation requires the *Dppa2/4* C-terminal domain, whereas the SAP domain is needed for later stages of embryonic development, reminiscent of the differential expression pattern of its murine orthologue (Siegel et al. 2009).

DPPA2 and DPPA4 control both *Dux* and LINE-1 transcription in mESCs. Interestingly, it was recently demonstrated that LINE-1 transcripts are necessary for TRIM28/Nucleolin-mediated repression of *Dux* in mESCs and in pre-implantation embryos (Percharde et al. 2018). Our results suggest that DPPA2 and DPPA4 influence a negative feedback loop involving *Dux* and its transcriptional repressors, which might explain how *Dux* is expressed as a brief pulse in pre-implantation embryos and in cycling 2C-like mESCs.

How DPPA2 and DPPA4 activate their targets remains to be formally determined. A high enrichment in heterochromatin marks, including DNA methylation and histone 3 lysine 9 dimethylation (H3K9me2), is found at promoters of genes regulated by DPPA4 in mESCs depleted for this factor (Madan et al. 2009). Moreover, general loss of the repressive chromatin mark histone 3 lysine 9 trimethylation (H3K9me3) was detected in mouse embryonic fibroblasts (MEFs) reprogrammed to induced pluripotent stem cells (iPSCs) when DPPA2 and DPPA4 were ectopically overexpressed (Hernandez et al. 2018). This suggests that DPPA2 and DPPA4 may bind and remodel the chromatin at the promoter of target genes to create an active environment and prevent the recruitment of repressor complexes.

## Material and Methods

### Cell lines and tissue culture

J1 mESCs (ATCC) were cultured in feeder-free conditions on 0.1% gelatin-coated tissue culture plates in Dulbecco’s modified Eagle’s medium (Sigma) containing 15% Fetal Bovine Serum (FBS, HyClone, Fisher Scientific) and supplemented with GlutaMAX (GIBCO), nonessential amino acids (Sigma), 2-mercaptoethanol (GIBCO), and 1,000 U/ml leukemia inhibitory factor (LIF, Millipore). WT and DUX KO E14 mESCs containing the MERVL regulatory sequence driving expression of a 3XturboGFP-PEST (Ishiuchi et al. 2015) were cultured on 0.1% gelatin-coated tissue culture plates in 2i medium (De Iaco et al. 2017). 293T cells were maintained in DMEM supplemented with 10% FCS. All cells were regularly checked for the absence of mycoplasma contamination.

### Plasmids and lentiviral vectors

Three single guide RNAs (sgRNAs) targeting sequences flanking *Dppa2* and *Dppa4* (Figure 1A) were cloned into px459 (version 2) using a standard protocol (De Iaco et al. 2017). Table S1 shows the primers used to clone the sgRNAs. The pLKO.1-puromycin shRNA vectors for *Dppa2* and *Dppa4* knockdown were ordered from Sigma (TRCN0000174599, TRCN0000175923, TRCN0000329372, TRCN0000329374) (De Iaco et al. 2017). *Dppa2* and *Dppa4* cDNAs were cloned from the genome of J1 mESCs and *Dppa4* ΔSAP from the genome of B62F1 blastocysts in a pDONR221 without STOP codon. The ORFs were then shuttled in a pTRE-3HA, which produces proteins with three C-terminal HA tags. The cDNAs, including *Dux*, followed by HA tags were finally cloned into a pWPTs-GFP HIV1-based transfer vector in place of the GFP reporter using the In-Fusion® HD Cloning Kit (Clontech) and the primers shown in Table S1. pMD2-G encodes the vesicular stomatitis virus G protein (VSV-G). The minimal HIV-1 packaging plasmid 8.9NdSB carrying a double mutation in the capsid protein (P90A/A92E) was used to achieve higher transduction of the lowly permissive mESCs (De Iaco et al. 2013).

### Production of lentiviral vectors, transduction and transfection of mammalian cells

Lentiviral vectors were produced by transfection of 293T cells using Polyethylenimine (PEI) (Sigma, Inc) (De Iaco et al. 2013). To generate stable KDs, mESCs were transduced with empty pLKO.1 vector or vectors containing the shRNA targeting *Dppa2* or *Dppa4* transcripts. Cells were selected with 0.4 μg/ml puromycin starting one day after transduction. To express HA-tagged DPPA2 or DPPA4, mESCs were transduced with the pWPTs-DPPA2 or pWPTs-DPPA4 plasmids and no selection was applied.

### Creation of KO mESC lines

J1 mESCs were co-transfected with px459 plasmids encoding for Cas9, the appropriate sgRNAs and puromycin resistance cassette by nucleofection (Amaxa™ P3 Primary Cell 4D-Nucleofector™ X Kit). 24 hours later, the cells were selected for 48 hours with 0.4 μg/ml puromycin, single-cell cloned by serial dilution, expanded and their DNA was extracted to detect the presence of WT and/or KO alleles. Three WT, three homozygous *Dppa2* KO and three homozygous *Dppa4* KO clones were selected and used in this study.

### Western blot

Actinβ (Abcam), DPPA2 (Millipore) and DPPA4 (Thermofisher) antibodies were used for Western blots.

### Fluorescence-activated cell sorting (FACS)

FACS analysis was performed with a BD FACScan system. J1 mESCs containing the MERVL-GFP reporter were subjected to FACS sorting with AriaII (BD Biosciences).

### Standard PCR, RT-PCR and RNA sequencing

For the genotyping of *Dppa2* and *Dppa4* WT and KO alleles, genomic DNA was extracted with DNeasy Blood & Tissue Kits (QIAGEN) and the specific PCR products were amplified using PCR Master Mix 2X (Thermo Scientific) combined with the appropriate primers (design in Figure 1A; primer sequences in Table S2).

Total RNA from cell lines was isolated using the High Pure RNA Isolation Kit (Roche). cDNA was prepared with SuperScript II reverse transcriptase (Invitrogen). Primers listed in Supplementary Table S1 were used for SYBR green qPCR (Applied Biosystems). Library preparation and 75-base-pair paired-end RNA-seq were performed using standard Illumina procedures for the NextSeq 500 platform. RNA-seq data generated in this study are available on GEO.

### GC-content analysis

GC-content of DPPA2, DPPA4 and DUX peaks was done first, by converting bed files to fasta with bedtools suite and then by using a home-made perl script to count DNA bases. For measurement of GC-content in DUX gene, a 200bp sliding window was used.

### ChIP and ChIP sequencing

ChIP and library preparation were performed as described previously (De Iaco et al. 2017). DPPA2-HA and DPPA4-HA ChIP was done using the anti-HA.11 (Covance) antibody. Sequencing was performed with Illumina NextSeq 500 in 75-bp paired-end reads run. ChIP-seq data generated in this study are available on GEO.

### RNA-seq datasets processing

RNA-Seq of Dppa2/4 KO vs WT and *Dux* KO vs WT mESCs (De Iaco et al. 2017) was mapped to mm9 genome using hisat2 aligner (Kim et al. 2015) for stranded and paired-end reads with options -k 5 --rna-strandness RF --seed 42 -p 7. Counts on genes and TEs were generated using featureCounts (Liao et al. 2014) with options -p -s 2 -T 4 -t exon -g gene_id -Q 10, using a gtf file containing both genes and TEs to avoid ambiguity when assigning reads. For repetitive sequences, an in-house curated version of the mm9 *open-3.2.8* version of *Repeatmasker* database was used (fragmented LTR and internal segments belonging to a single integrant were merged) (Ecco et al. 2016). Single-cell RNA-Seq of 2C-like cells (E-MTAB-5058) datasets was downloaded from GEO (Eckersley-Maslin et al. 2016). The processing of the single-cell followed a previously published pipeline. Single-cell RNA-Seq mouse early embryo development data were reanalyzed from (De Iaco et al. 2017)

### RNA-seq analysis

Normalization for sequencing depth and differential gene expression analysis was performed using Voom as it has been implemented in the limma package of Bioconductor, with total number of reads on genes as size factor (De Iaco et al. 2017). TEs overlapping exons or having less than 1 read per sample in average were removed from the analysis. To compute total number of reads per TE family/subfamily, counts on all integrants were summed up using multi-mapping read counts with fractions (featureCounts with options -M --fraction - p -s 2 -T 4 -t exon -g gene_id -Q 0) to compensate for potential bias in repetitive elements.

### ChIP-seq data processing

ChIP-seq dataset of DUX overexpressed in mESCs (GSE85632) was downloaded from GEO (Hendrickson et al. 2017). Reads were mapped to the mouse genome assembly mm9 using Bowtie2 using the sensitive-local mode. MACS2 (the exact parameters are: macs2 callpeak -t $chipbam -c $tibam -f BAM -g $org -n $name -B -q 0.01 --format BAMPE) was used to call peaks (De Iaco et al. 2017). To compute the percentage of bound TE integrants in each family, we used bedtools suite.

### Methodology for statistics

Enrichment of TE subfamilies were done using hypergeometric tests, comparing the number of peaks having at least 50% overlap with TEs to the expected number. Enrichment of peaks around TSS also used a hypergeometric test to compare the occurrence of peaks in a subset of TSS to the expected. When the expression data was compared between two conditions, a paired student t-test was used. To correct for different sample sizes, we used 1000 subsampling permutations using the smallest sample size to get the median p-val. For comparing enrichment between Dppa2 and Dppa4 down-regulated genes, a gene-set enrichment analysis was used with *phenoTest* library from *Bioconductor* with default options.

### Coverage plots

Raw ChIP-seq data were mapped on an index containing only the *Dux* gene using the same parameters as described in the methods when mapping on the genome. ChIP-seq signals on the locus were extracted from the bigWigs and normalized for sequencing depth (reads per hundred millions when mapped on genome) using the pyBigWig python library. Replicate signals were averaged and then smoothed using a running average of window 250bp prior to plotting.

### MSAplot

Fasta sequences for the TE familes of interest were extracted from the mm9 genome assembly and aligned using the mafft aligner with parameters: --auto –reorder. Regions in the alignment consisting of more than 95% of gaps were trimmed out. For each integrant in the family, the ChIP-seq signal was extracted from the bam, scaled to the interval [0, 1] and plotted on top of the alignment using python. Finally, the average of the signals was plotted on top of the MSAplot.

## Acknowledgements

We thank the Gene Expression and Flow Cytometry Core Facilities (EPFL) for technical assistance. This work was financed through grants from the Swiss National Science Foundation, the Gebert-Rüf Foundation, FP7 MC-ITN INGENIUM (290123), and the European Research Council (ERC 694658) to D.T.

## Author contributions

A.D.I and D.T. conceived the project, designed the experiments, analyzed the data and wrote the manuscript; A.D.I. carried out the experiments; A.C. and J.D. performed the bioinformatics and statistical analyses.

